# Deficiency in *Lyst* function leads to accumulation of secreted proteases and predisposition to mechanic stress-induced retinal detachment

**DOI:** 10.1101/2021.07.23.453557

**Authors:** Xiaojie Ji, Lihong Zhao, Ankita Umapathy, Bernard Fitzmaurice, Jieping Wang, David S. Williams, Bo Chang, Jürgen K. Naggert, Patsy M. Nishina

## Abstract

Chediak–Higashi syndrome, caused by mutations in the <u>Lys</u>osome <u>T</u>rafficking Regulator (*Lyst*) gene, is a recessive hypopigmentation disorder characterized by albinism, neuropathies, neurodegeneration, and defective immune responses, with enlargement of lysosomes and lysosome-related organelles. Although recent studies have suggested that *Lyst* mutations impair the regulation of sizes of lysosome and lysosome-related organelle, the underlying pathogenic mechanism of Chediak–Higashi syndrome is still unclear. Here we show striking evidence that deficiency in LYST protein function leads to accumulation of photoreceptor outer segment phagosomes in retinal pigment epithelial cells, and reduces adhesion between photoreceptor outer segment and retinal pigment epithelial cells in a mouse model of Chediak–Higashi syndrome. In addition, we observe elevated levels of cathepsins, matrix metallopeptidase (MMP) 3 and oxidative stress markers in the retinal pigment epithelium of *Lyst* mutants. Previous reports showed that impaired degradation of photoreceptor outer segment phagosomes causes elevated oxidative stress, which could consequently lead to increases of cysteine cathepsins and MMPs in the extracellular matrix. Taken together, we conclude that the loss of LYST function causes accumulation of phagosomes in the retinal pigment epithelium and elevation of several extracellular matrix-remodeling proteases through oxidative stress, which may, in turn, reduce retinal adhesion. Our work reveals previously unreported pathogenic events in the retinal pigment epithelium caused by *Lyst* deficiency, which may place Chediak–Higashi syndrome patients at increased risk for retinal detachment. The same pathogenic events may be conserved in other professional phagocytic cells, such as macrophages in the immune system, contributing to overall Chediak–Higashi syndrome pathology.

## INTRODUCTION

Chediak–Higashi syndrome (CHS) is a rare autosomal recessive disease characterized by albinism of the skin and hair, as well as hypopigmentation of the eye and additional eye pathologies including photophobia and macular hypoplasia associated with decreased visual acuity [1–4]. Patients also display progressive neurologic dysfunction, including motor and sensory neuropathies, ataxia, and progressive neurodegeneration [1, 2, 5–9]. The most detrimental pathology is, however, recurrent bacterial infections, which predominantly affect the respiratory tract, skin, and mucous membranes. These infections are due to the dysfunction of polymorphonuclear leukocytes [1, 2, 10, 11]. The majority of CHS cases progress to a life-threatening lymphoproliferative accelerated phase characterized by massive hemophagocytic lymphohistiocytosis after exposure to Epstein-Barr virus [2]. Antibiotic treatments and hematopoietic stem cell transplantation have been used to combat recurrent infections and immunological complications; but these treatments target the symptoms, not the underlying pathogenic mechanism(s) [1, 2, 12].

CHS is caused by mutations in the ubiquitously expressed <u>Lys</u>osome <u>T</u>rafficking Regulator (*Lyst*) gene, which encodes LYST, a <u>Be</u>ige <u>a</u>nd <u>C</u>hediak-<u>H</u>igashi (BEACH) domain-containing protein [12–15]. The loss of LYST function results in enlarged lysosomes and lysosome-related organelles (LROs) in all cell types examined [1, 14, 16–22]. Functional studies using several model organisms have previously led to two distinct hypotheses for the effects of LYST in the regulation of LRO sizes: LYST may restrict homotypic lysosome fusion [23–28] by inhibiting membrane docking and fusion [23], or alternatively, LYST may promote lysosome fission [17, 29, 30]. However, a recent report indicates that LYST function is likely to be far more complex than a simple role in either lysosomal fusion or fission, and suggests that LYST may regulate fusion through fission-mediated recycling of the fusion machinery during lysosomal maturation [31]. Despite years of research, the exact molecular function of LYST remains unclear. Given that LYST is an extremely large protein, approximately 430 kDa in size, and contains multiple WD40 domains implicated in protein-protein interaction, it is likely that LYST has many alternate functions that are dictated by its interaction with binding partners. Thus, loss of LYST function may cause various cellular defects that have yet to be elucidated.

In this study, we examined the downstream effects of LYST dysfunction on the cellular pathology of the retinal pigment epithelium (RPE). The RPE is a monolayer of post-mitotic polarized epithelial cells, situated between the photoreceptors and the choroid; it is the primary caretaker of photoreceptor health and function [32]. One of the primary functions of the RPE is to engulf and degrade the distal tips of photoreceptor outer segments (POSs) [33, 34]. Because each RPE cell serves many photoreceptor cells (200 in the mouse central retina) [35], they are tasked with degrading extraordinary amounts of POS material on a daily cycle [36, 37].

By taking advantage of a novel mutant mouse strain bearing a mutation in *Lyst, Lyst^bg-18J^*. We found that, in addition to enlarged and redistributed lysosomes and accumulation of phagosomes in RPE cells, there is reduced adhesion between the RPE and the neural retina, causing higher risk in retinal detachment. We show that the accumulation of phagosomes is associated with increased oxidative stress and an elevation of a group of proteases, including cathepsins B, L, and S, and matrix metalloprotease 3 (MMP3), which are likely secreted into the interphotoreceptor matrix (IPM) and may contribute to the retinal adhesion defect. The disturbances in lysosomal enzymatic activities may have a profound impact on tissue integrity beyond the RPE. The same mechanism is likely to also exist in the immune system, where elevation of secreted proteases cleave cell surface proteins potentially leads to the reduced immune response observed in CHS [38]. Our results establish a series of events for the pathogenic mechanism of CHS and suggests an important new entry point for therapeutic intervention.

## MATERIALS AND METHODS

### Mice

The *Lyst* mutant *bg-18* (*Lyst^bg-18J^*) was first identified by the JAX Mouse Mutant Resource as a spontaneous mutation, *nm2144*. Experimental animals were housed in the same mouse room and under the same 14-hr light / 10-hr dark cycle from birth. The light cycle was 6AM to 8PM. All experiments were approved by the Institutional Animal Care and Use Committee and conducted in accordance with the ARVO Statement for the Use of Animals in Ophthalmic and Vision Research.

### Indirect Ophthalmoscopy, Optical Coherence Tomography and Electroretinography

Indirect ophthalmoscopy, optical coherence tomography (OCT) and electroretinography (ERG) were performed as described [39]. For indirect ophthalmoscopy and OCT, 11 10-week-old wild type C57BL/6J male and 2 female mice, and 4 17-week-old mutant male mice, and 11 19-month-old male mice were used. Three wild type and 3 mutant 2-month-old male mice were used for ERG.

### RNA preparation and reverse transcription

After carbon dioxide (CO_2_)-induced euthanasia, mouse eyes were dissected in DEPC-treated water. The posterior eyecup was separated from connective tissues and the iris epithelium, cornea, and lens. For RPE only RNA preparations, the RPE was peeled from the neuroretina. RPE from both eyes of each animal was pooled. Poly A+ RNA from the RPE was extracted using Dynabead mRNA DIRECT Micro Kit (Invitrogen) according to the manufacturer’s instructions. RNA concentration was quantified using a NanoDrop ND-1000 Spectrophotometer (NanoDrop Technologies). cDNA was synthesized using the RETROscript Kit (ThermoFisher Scientific).

### Quantitative real-time PCR

Real-time PCR was performed using Bio-Rad iTaq mixture on Bio-Rad iCycler 96 thermocycler equipped with a CCD image detector, using protocols with a melt curve analysis. Only primers generating a solid major peak without obvious minor peaks in the melting curve were used. Samples were collected from three wild type and three homozygous *bg-18* mice. Each sample was subjected to three technical replications. Primers, PCR procedure and data analysis are described in detail in Supporting Information. PCR products were resolved by Metaphor agarose gel electrophoresis to confirm the expected sizes.

### Western blot analysis

After mice were euthanized by CO_2_ asphyxiation, the eyes were dissected in ice-cold 1X Phosphate-Buffered Saline (PBS). Individual eyecups or pooled RPE from the same animal were homogenized in ice-cold RIPA buffer [50 mM Tris.HCl pH 8.0, 1 mM EDTA, 150 mM NaCl, 1% NP-40, 0.1% SDS, and 0.5% sodium deoxycholate], freshly supplemented with phosphatase inhibitors and protease inhibitors [100 mM sodium orthovanadate, 10 mM ammonium molybdate, 0.2M sodium pyrophosphate, 1 M sodium fluoride, 0.1 M phenylmethanesulfonyl fluoride and 1% of protease inhibitor cocktails (EMD Millipore)]. Insoluble material from the lysate was removed by centrifugation for 10 minutes at 10,000 x g at 4°C. Protein concentration was measured with Assay-free Card (EMD Millipore) and Bradford Ultra (Expedeon) kits. Equal amounts of total protein representing ~10% of the whole eyecup were used for western blot analysis as described [39]. Results were quantified using Fiji software (https://fiji.sc/).

Antibodies against phospho-MERTK (FabGennix, PMKT-140AP, rabbit polyclonal, 1:750), 4-HNE (Abcam, ab46545, rabbit polyclonal, 1:500), Glial Fibrillary Acidic Protein (GFAP, DAKO, Z0334, rabbit polyclonal, 1:1000), glyceraldehyde 3-phosphate dehydrogenase (GAPDH, Cell Signaling, 2118, rabbit monoclonal, 1:1000), phospho-ERM (Cell Signaling, 3726T, rabbit monoclonal, 1:1000), cathepsin B (Cell Signaling, 31718T, rabbit monoclonal, 1:1000), and α-tubulin (Santa Cruz, sc-53030, rat monoclonal, 1:1000) were used.

### Quantification of melanin

After lysis of tissue samples with RIPA buffer, melanosomes precipitate in the insoluble pellet. The insoluble pellet was dissolved in 1 N NaOH at 80°C for 2 hr. Absorbance of commercial melanin pigment (Sigma) at defined concentrations was measured at 405nm to establish a standard curve. The melanin content of each sample was assessed at O.D. 405 and compared to the standard curve. The pigment concentration was normalized to the total protein concentration within each sample (μg melanin / mg protein).

### Retinal adhesion assay

Retinal adhesion assays were performed at two time points, 9:00 AM (3 hours after onset of the light cycle) and 3:00 PM (9 hours after onset of the light cycle). Three wild type and three mutant 7-week-old mice were used for each time point. Enucleated eyes were submerged in 20 mM HEPES-buffered Hanks’ saline solution containing calcium and magnesium (Mediatech, Inc., A Corning Subsidiary) at room temperature to preserve retinal adhesion. Eyecups were dissected as described above. A single radial cut toward the optic nerve was made to flatten the eyecups. The neural retina was slowly peeled from the underlying RPE and sclera with forceps from one side of the cut edge to the other. The peeled-off neural retina was then flattened, facing upward on a glass slide for imaging.

The imaging results were confirmed with western blot analysis, which measured the amount of RPE-specific proteins in the peeled-off neural retina. Blots were incubated with primary antibodies against ezrin (Cell Signaling, 3145, rabbit polyclonal, 1:1000), GFAP, and GAPDH. Ezrin, a marker for RPE apical microvilli, was used to detect the amount of RPE attached to neural retina after separation from the RPE. GFAP and GAPDH were used as loading controls.

### Transmission electron microscopy

Mice were perfused intracardially with buffered 1.2% (wt/vol) paraformaldehyde and 0.8% glutaraldehyde, and eyecups were processed using a standard procedure as described previously [40]. The eyes for ultrastructural analyses were fixed in an ice-cold fixative solution for 3 h. The anterior segment was removed and the posterior segment cut into 1 × 2 mm blocks. Additional fixation with 0.25% glutaraldehyde/0.2% paraform- aldehyde fixative was performed for 2–8 h followed by post-fixation with 1% osmium tetroxide. The dehydrated blocks were embedded in plastic. Tissue sections were cut and stained with uranyl acetate and lead citrate and examined with a JEM-1230 transmission electron microscope (JEOL, Ltd).

### Immunohistochemistry (IHC)

After mice were sacrificed by CO_2_ asphyxiation, eyes were enucleated and submerged in ice-cold 4% paraformaldehyde (PFA) in PBS or acetic acid/methanol/PBS (1:3:4) overnight. Standard IHC was conducted as described [39]. Antibodies against rhodopsin (RET-P1; Thermo Fisher Scientific, MS-1233-R7, dilute each drop to 250 μL), ezrin (Cell Signaling, 3145, rabbit polyclonal, 1:200), ezrin (Santa Cruz, sc-6409, 1:200), and cathepsin B (Cell Signaling, 31718T, rabbit monoclonal, 1:200) were used. These antibodies detected bands of expected or reported sizes on western blot analyses.

### Immunostaining of RPE flat mounts

Enucleated eyes were submerged in ice-cold 1X PBS and dissected as described above. The neural retina was separated from the RPE-choroid-sclera. The isolated RPE-choroid-sclera was fixed for 30 minutes in 4% PFA. Subsequently, the tissue was washed three times with 1X PBS, permeabilized with PBST (PBS with 0.1% Tween-20) for 20 minutes, and blocked with 5% donkey serum in PBST for 1 hr. Tissues were incubated with primary antibodies against rhodopsin (RET-P1; Thermo Fisher Scientific, MS-1233-R7, dilute each drop to 250 μL) and ZO-1 (Life Technologies, 61-7300, 1:200) in blocking solution overnight at 4°C, washed with PBST, and incubated with fluorophore-conjugated secondary antibodies for 2 hr at room temperature. Nuclei were counterstained by incubating with DAPI for 20 minutes. The whole RPE-choroid-sclera tissue was then flat mounted, with the RPE side up, onto slides and with coverslips for imaging. Fluorescent images were captured with a Leica SP5 confocal microscope (Leica Microsystems), and identical imaging parameters were applied to both wild type and *Lyst^bg-18^* mutant RPE flat mounts.

For image analysis, only rhodopsin-positive POS fragments with diameter 0.5 μm-2.5 μm were counted by using the Imaris 9.1 software (Bitplane USA, Concord, MA, USA), with the aid of the ‘Surfaces’ rendering tool. Clustered POS fragments whose boundaries were difficult to delineate or those that clearly appeared to be cell surface adherent POS were not included in counts. The results were quantified and analyzed using unpaired *t*-test with Prism 6 software.

## RESULTS

### Pathogenic effects of *Lyst* deficiency in the RPE

We have identified a new mutant mouse *bg-18* (B6.Cg-*Lyst^bg-18J^*/Boc, JR#028230, The Jackson Laboratory) that recapitulates many of the pathologies observed in human CHS patients and resembles the *beige* (*Lyst^bg-J^*) mutant previously reported [22]. Homozygous *bg-18* mutant mice, like *beige* mutant mice, display a dark grey coat color on the C57BL/6J background, compared with wild type littermates (Fig. 1A). The underlying mutation was first inferred as a mutation in *Lyst* in a whole exome sequencing project on 91 mutant mice [41]. Subsequent genome mapping, complementation test with the *beige* mutant, and sequencing of *Lyst* cDNA and genomic DNA from the *bg-18* mutant definitively placed the causal mutation in the *Lyst* locus, as an intronic mutation causing a skipping of exon 10 resulting in a frameshift of the mRNA and truncation of the encoded LYST protein (Fig. S1). Details describing the genetic studies are included in the Supporting Text. We will refer to the *bg-18J* mutants as *Lyst* mutant mice, hereafter. This mutant was used to study the pathogenic effects of *Lyst* deficiency in this report.

**Fig. 1.**
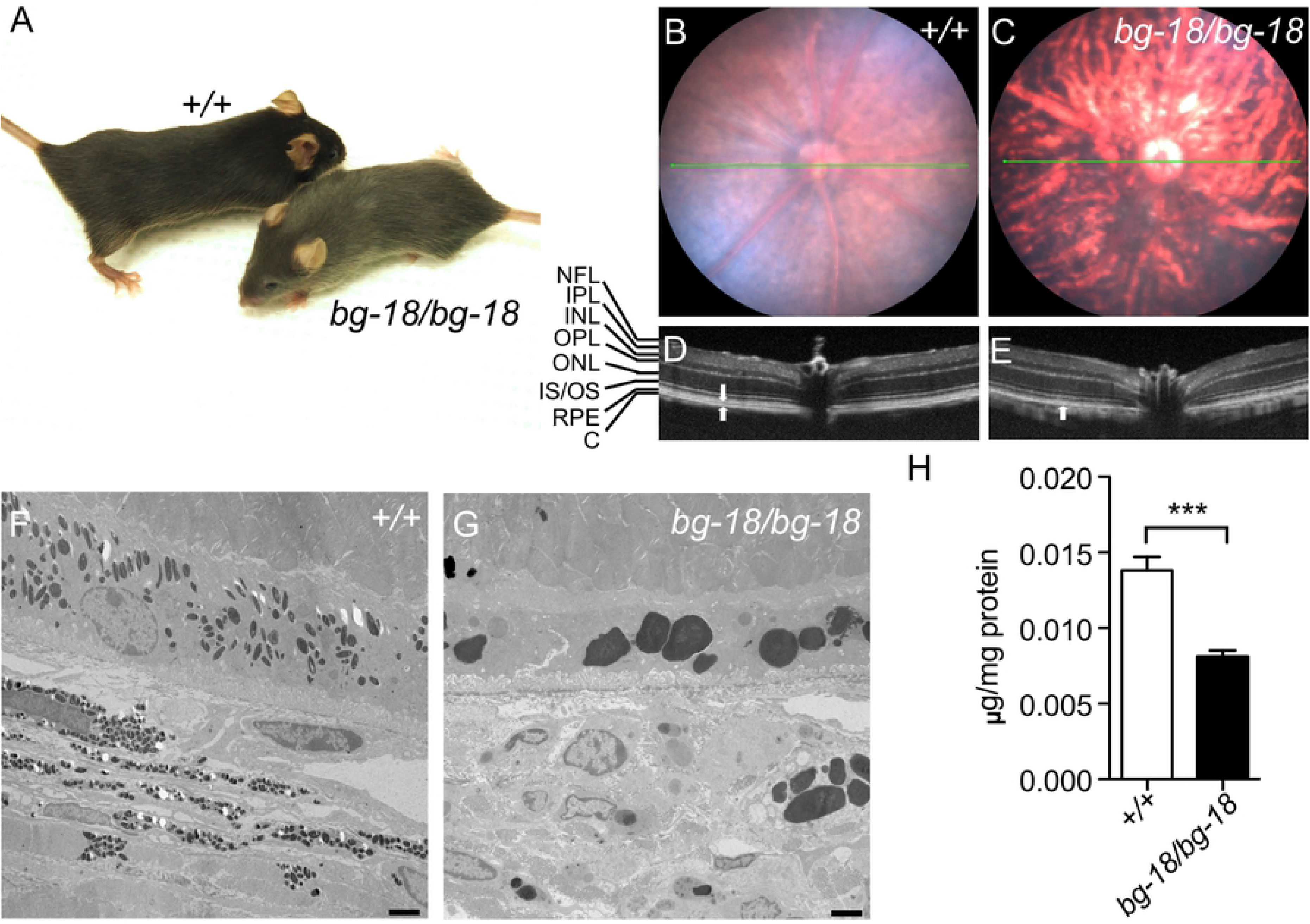
The clinical and pathological effects of the *bg-18* mutation on mouse retina. **A.** 5-week-old wild type (+/+) and *bg-18/bg-18* mutant littermates on the C57BL/6J background. Note the lighter coat color of the *bg-18* mutant. **B-E.** Abnormalities of pigmentation and reflective properties in the *bg-18* mutant retina. Fundus (B, C) and OCT (D, E) images of wild type (B, D) and *bg-18* mutant (C, E) retinas from 4-month-old mice. Retinal layers are labeled to the left. NFL: neurofilament layer; IPL: inner plexiform layer; INL: inner nuclear layer; OPL: outer plexiform layer; ONL: outer nuclear layer; IS/OS: inner segment/outer segment; RPE: retina pigment epithelium; C: choroid. Note that there are two hyper-reflective layers in the wild type retina at the location of RPE and choroid (arrows), whereas only one layer is detected in the mutant retina (arrow). **F-G.** Electron micrographs of retinas from the 11-week-old wild type (F) and *bg-18* mutant (G) retinas. Scale bar = 2μm. Three mice of each genotype were examined. **H.** Melanin level is reduced in the *bg-18* mutant retina from 5-week-old mice. ***: p<0.001, mean ± SD, n=3.

The fundus images of homozygous *Lyst* mutants show an uneven distribution of retinal pigmentation compared to those of wild type mice (Fig. 1B, C). Optical coherence tomography (OCT) reveals two hyper-reflective layers corresponding to the RPE and choroid in the wild type retina, while only one layer is observed at the same location in the mutant retina (Fig. 1D, E), suggesting an alteration of the posterior retina.

By using transmission electron microscopy (TEM), we found that melanosomes in the RPE were remarkably enlarged both in the RPE and choroid of the *Lyst* mutants, relative to melanosomes in controls (Fig. 1F, G). Because the melanosomes tended to aggregate without distinct boundaries (Fig. S2), we were unable to distinguish individual organelles and quantitate their number. However, a significant reduction of melanin concentration was observed in the *Lyst* mutant eyecups compared with wild type controls (Fig. 1H), indicating a reduction in melanogenesis or an increase in melanin turnover.

At 2 months of age, *Lyst* mutants show similar rod and cone electroretinography (ERG) responses to wild type controls (Fig. S3), indicating no obvious electrophysiological defects or retinal degeneration at this age. ERG measurements were also attempted on aged *Lyst* mutants but were not successful as their eyes could not be dilated. However, retinal degeneration was not observed at 19 months of age in OCT images of *Lyst* mutant eyes (Fig. S4).

### Increased number of phagosomes in the *Lyst* mutant RPE

The tips of rod OS are phagocytosed by the RPE immediately after light onset [36, 37]. We counted the numbers of phagosomes containing rod outer segments (OS) in RPE flat mounts from wild type and *Lyst* mutant mice at 7 to 8 weeks of age. At the time of light onset, the number of RPE phagosomes in *Lyst* mutant RPE were slightly higher compared with wild type control (Fig. 2A, D). We also compared the numbers of RPE rhodopsin-positive phagosomes in IHC images of 5-week-old wild type and *Lyst* mutant eyes at 0.5, 1, and 2 hr after light onset and found that the number of phagosomes were higher in the mutant RPE at all time points examined (Fig. S5). The most remarkable differences in the number of rhodopsin-positive phagosomes in *Lyst* mutant RPE cells when compared to wild type, however, was at 3 or 9 hr after light onset (Fig. 2B, C, E, F, G; Fig. S6).

**Fig. 2.**
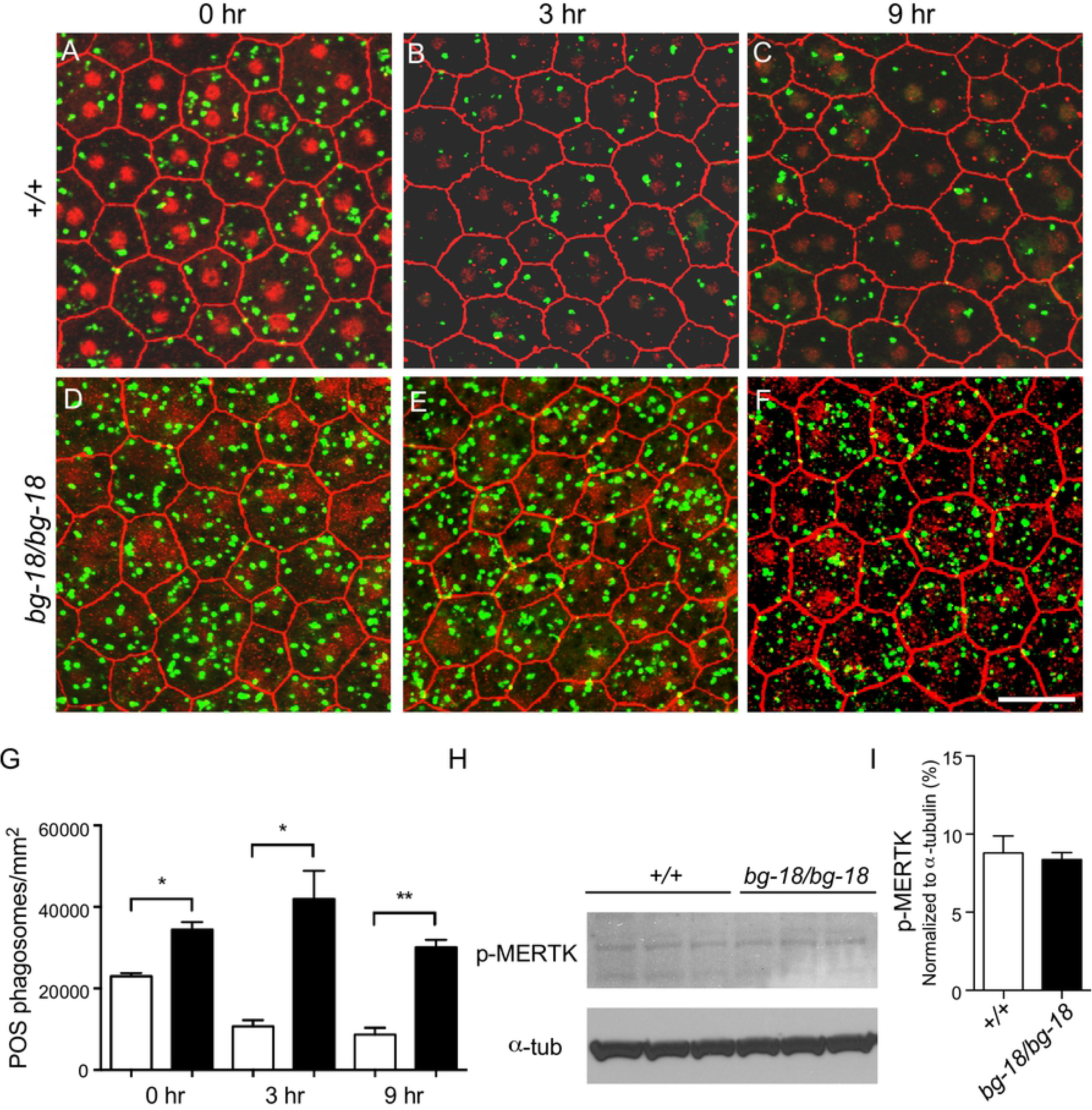
Accumulation of phagosomes in *bg-18* RPE cells. **A-G.** Number of phagosomes in RPE from 7 to 8-week-old wild-type and *bg-18* mutant mice. Wild type (A-C) and *bg-18* mutant (D-F) RPE was incubated with antibodies against ZO-1 (red) and rhodopsin (green), after dissection at 0 (A, D), 3 (B, E), and 9 (C, F) hours after the onset of light at 6 AM. Note that the ZO-1 staining in the nuclei is an artifact of over-staining with DAPI which can result in bleed over signal into the red channel. This does not affect the green channel. Scale bar = 50μm. Images covering larger areas are shown in Fig. S6. The results were quantified and analyzed using unpaired *t*-test (G). *: P<0.05, **: P<0.01, mean ± SD, n=3. **H-L.** Phagocytosis is not affected in the *bg-18* mutant RPE from 5-week-old mice, as assessed by the phospho-MERTK level using western blot analysis (H). Quantified results are shown in (I) as mean ± SD, n=3.

The increase in RPE phagosomes may either be due to an elevation of phagocytic activity or reduction in lysosomal degradation of engulfed phagosomes in the RPE. To differentiate between these two possibilities, the level of active, phosphorylated MER proto-oncogene tyrosine kinase (MERTK), which has been reported to increase upon the onset of phagocytosis [42], was assessed by western blot analysis. No obvious change was observed right after light onset (Fig. 2H, I), the time point of peak phagocytosis reported previously [43]. As MERTK is only localized on the apical surface of the RPE cells, this result indicates that phagocytic activity is not affected in the RPE of *Lyst* mutants. In addition, we found that the phosphorylated MERTK was still mainly observed in the *Lyst* mutant RPE at the time of lights-on but not at later sampled time points (Fig. S5). This finding indicates that phagocytosis of POS tips in LYST-deficient RPE cells is not prolonged and suggests that the phagocytic activity and the duration of phagocytosis are not affected by the *Lyst* mutation. Thus, we propose that the degradation of engulfed phagosomes, by lysosomes, is impaired, which results in the accumulation of undigested phagosomes in the RPE. Previous research has shown that the fusion between phagosomes and lysosomes is blocked by the re-distribution of lysosomes to perinuclear clusters induced by lipopolysaccharide (LPS) treatment in dendritic cells [44]. Other studies have demonstrated giant and perinuclear lysosomes in *Lyst*-deficient cells [19, 45, 46]. Our *Lyst^bg-18J^* mice recapitulate this subcellular phenotype of lysosomes (Fig. S7). We postulate that the perinuclear aggregation of lysosomes contributes to the accumulation of phagosomes in the RPE of *Lyst* mutants by blocking phago-lysosomal fusion.

### Reduced retinal adhesion in LYST deficient mice

When we performed histology, and compared the wild type (Fig. 3A-C) to the 3-week-old *Lyst* mutant retinas (Fig. 3D), we observed small areas of detachment of the neural retina from the RPE in sections from every homozygous *Lyst* mutant mouse examined (n>3 for each time cohort). The detachment was observed as early as 1 month of age (Fig. 3E), which appeared as larger areas of detachment by 3 months of age (Fig. 3F). However, retinal detachment is not observed by OCT at 1 or 3 months (Fig. S8). The detachment observed by histology may result from a meaningful artifact that occurs during enucleation of the eye and histological sample preparation due to the reduced retinal adhesion and is, therefore, not observed by OCT, which is an *in situ* technique. This suggests that the *Lyst* mutation results in the reduction of retinal adhesion, which only manifests as a full-blown detachment when the retina is stressed mechanically.

**Fig. 3.**
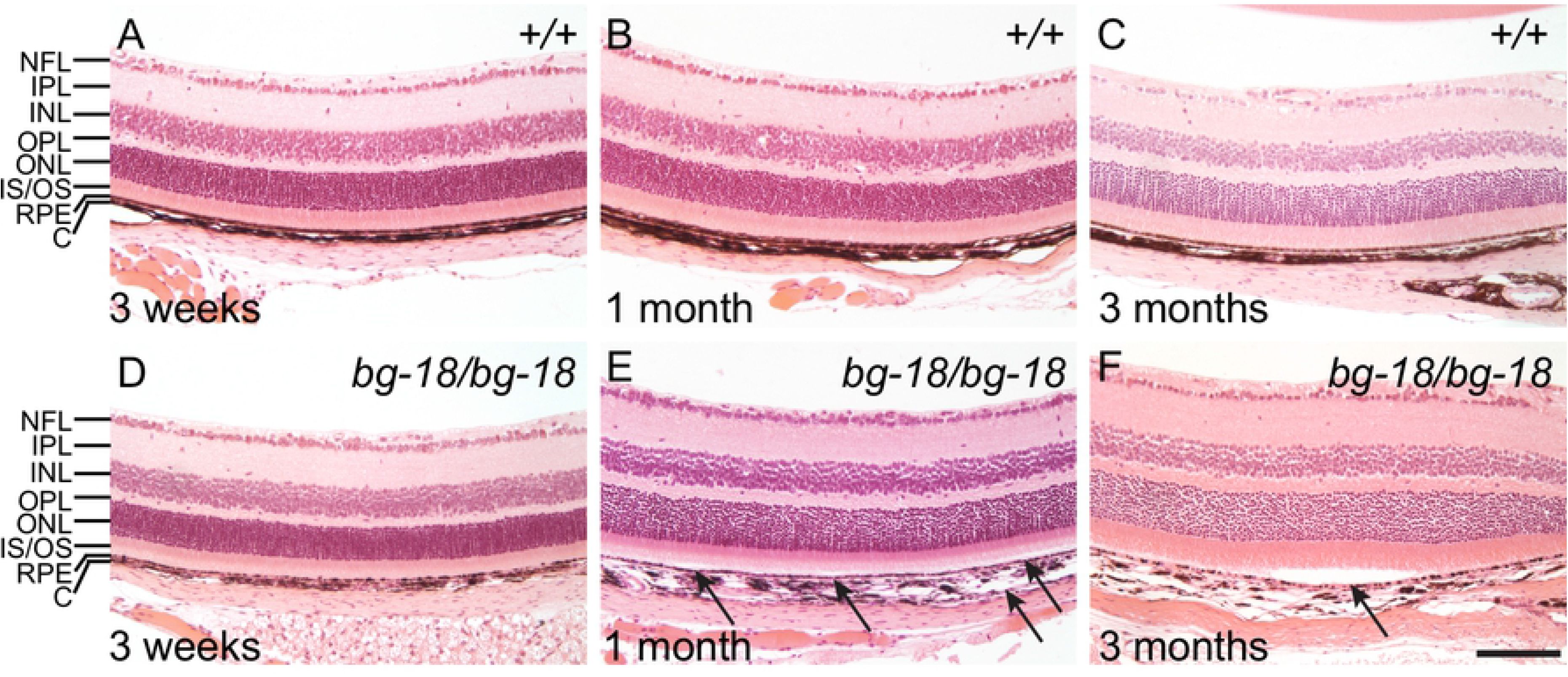
Progressive retinal detachment in *bg-18* mutant eyes as shown by H&E staining. Wild type (A-C) and mutant (D-F) retinas are compared at 3 weeks (A, D), 1 month (B, E), and 3 months (C, F) of age. Note the separation between the neural retina and RPE in the 1-month-old mutant eye (E) and the larger area of detachment in the retina of the 3-month-old mutant (F), marked by arrows. Retina layers are labeled as in Fig. 1D. Scale bar = 100μm.

Separation of the neural retina from its supportive structures can lead to significant visual defects [47–52]. The microvilli-rich apical domains of RPE cells, which form the outermost layer of the retina, interdigitate with the photoreceptor outer segments. Thus, apical domains of RPE cells or even entire RPE cells may remain attached when the neural retina is mechanically peeled from the RPE [42]. To determine if the retinal detachment observed by histology, but not by OCT, was an artifact of histological sample preparation or the consequence of reduced retinal adhesion insufficient to induce detectable retinal detachment *in vivo,* we compared the adhesion between the RPE and neuroretina in wild type and *Lyst* mutants accordingly. In the wild type peeled-off neural retina, RPE pigment was abundantly attached to the surface of photoreceptor outer segments (POS); whereas the peeled-off neural retina of the age-matched *Lyst* mutant mice (separated under the same experimental conditions and at similar times of day), were almost devoid of RPE pigment (Fig. 4A, B). To quantify the alteration in adhesion, the level of ezrin, an RPE microvillus marker was measured by western blot analysis in the peeled-off retinas. Compared to wild type neural retina, the ezrin levels were significantly reduced in the *Lyst* mutant peeled-off neural retina (Fig. 4C, D). In contrast, ezrin levels were similar in the wild type and mutant whole eyecups (Fig. 4E, F). Therefore, the reduction in ezrin levels in the mutant peeled-off neural retina is likely due to poor RPE-retina adhesion in the *Lyst* mutant mice, and not due to a decrease in ezrin levels in the RPE microvilli.

**Fig. 4.**
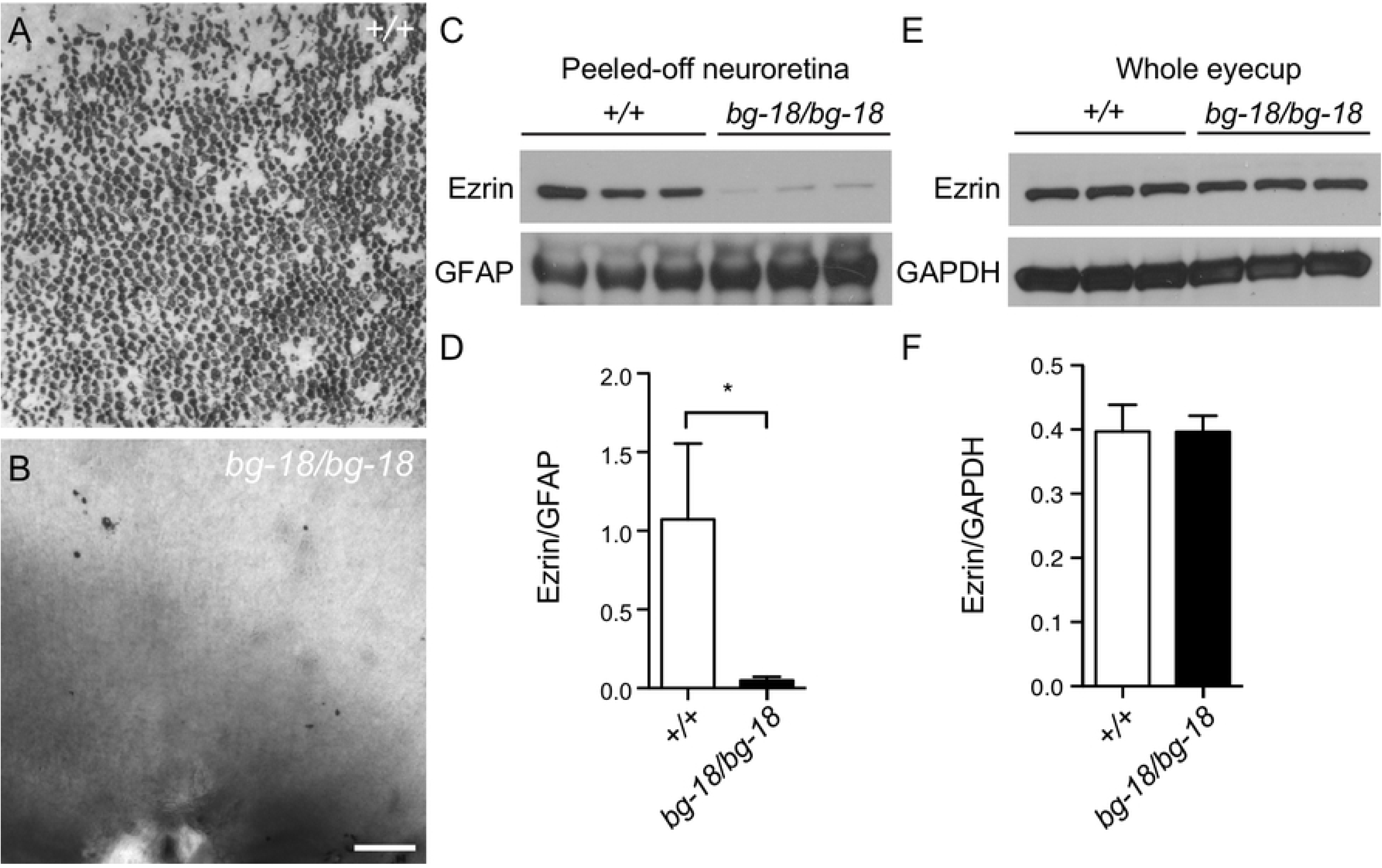
Reduced retinal adhesion in *Lyst* mutant mice. **A-B.** Pigment remaining on peeled-off retina from 7-week-old wild type (A) and mutant (B) eyes. Scale bar = 100μm. **C-D.** Reduced ezrin level in the mutant neuroretina shown by western blot analysis (C). Quantified results are shown (D). Ezrin levels were normalized to the levels of GFAP. **E-F.** Total ezrin levels are similar in mutant and wild type control eyecups in western blot analysis (E) and quantified results (F). *: P<0.05, mean ± SD, n=3.

### Levels of cysteine cathepsins and MMP3 are increased in the *Lyst* mutant eyecups

The POS and RPE microvilli of the *Lyst* mutant mice were not as densely packed as those in the wild type retina, but the structure of the processes were still clearly visible and comparable to that of the wild type mice (Fig. S9). In addition, we also examined ERM (ezrin, radixin, moesin) proteins that are activated upon phosphorylation (Thr567 of ezrin, Thr564 of radixin, Thr558 of moesin) and link membrane-associated proteins to actin filaments at the cell cortex, making them important for RPE microvilli morphology. No significant differences were observed in the level of phospho-ERM between wild type and *Lyst* eyecups (Fig. S9). These results suggest that the reduced adhesion between the neural retina and RPE in the mutant eye is unlikely to be due to structural alterations.

Adhesion of the neuroretina to the RPE is mediated by the interphotoreceptor matrix (IPM), a retina-specific type of extracellular matrix (ECM) between the RPE and POS. Previously, it was shown that proteases, such as matrix metalloproteinases (MMPs) and cysteine cathepsins, are found in the ECM and play important roles in ECM remodeling [53–56]. Quantitative real-time PCR showed up-regulation of transcripts of cathepsins and MMP12 in *Lyst* mutant ocular tissues[57]. To confirm previous reports and pinpoint the precise locations where the up-regulations actually occur, we tested transcripts of cathepsin B, L, and S and MMP3 in the RPE, because it appears to be the most affected layer in the retina by the *Lyst* mutation. Consistent with previous reports, we also found that the transcription levels of MMP3, and cathepsin B, L, and S assessed by quantitative real-time PCR, are increased in the RPE of the *Lyst* mutants relative to control (Fig. 5A-D). However, in both wild type and the *Lyst* mutant RPE, MMP3 and cathepsin S transcriptional expression was very low, and the protein levels, assessed by western blot analysis, were below detection level. Thus, in this study, our analysis focused on cathepsin B, as it is the most abundant cysteine protease in the RPE [58]. However, other aforementioned proteases upregulated by *Lyst* deficiency may play similar roles in the IPM.

**Fig. 5.**
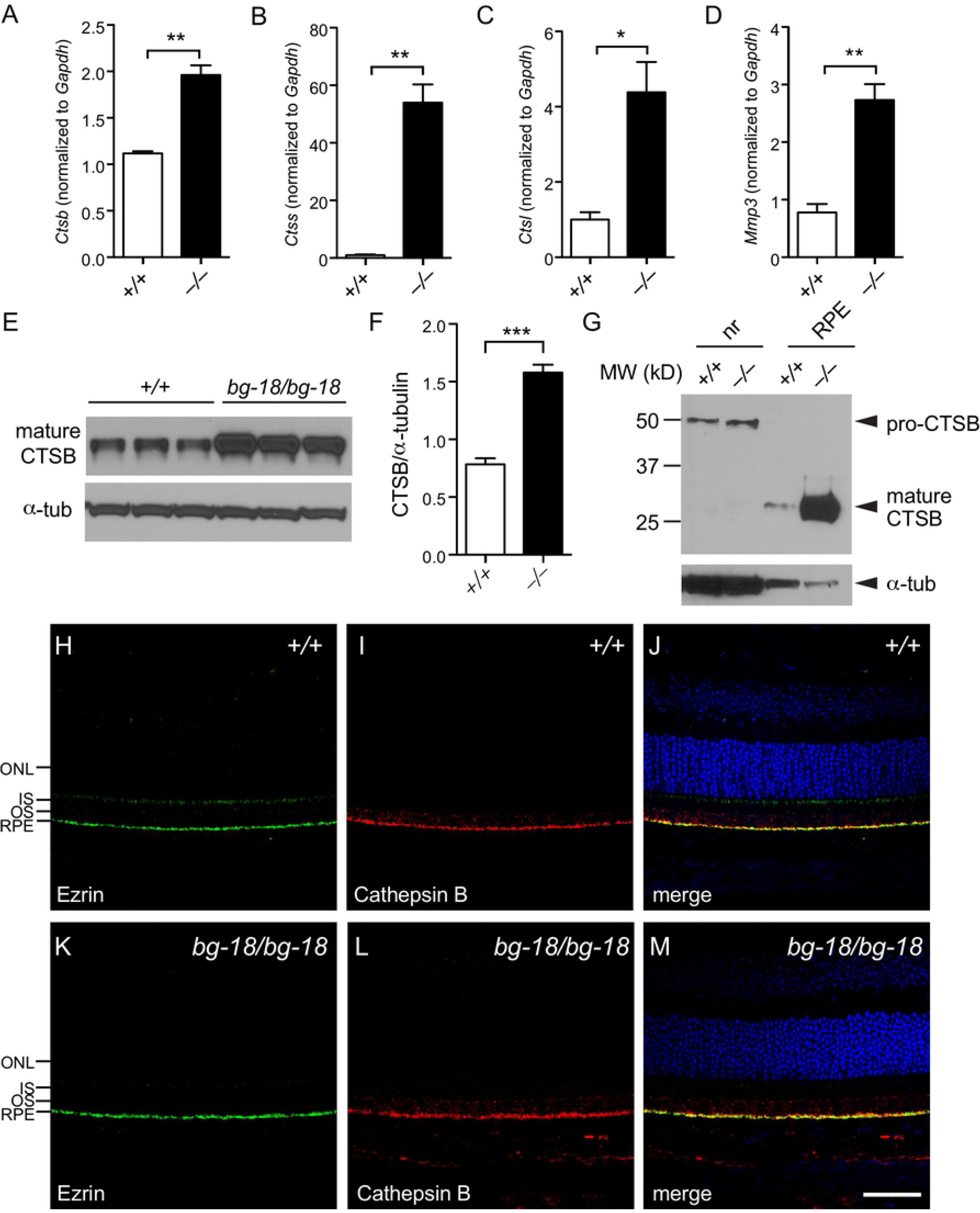
Elevation of cathepsins and MMP3 in the *Lyst* mutant RPE. **A-D.** Quantitative RT-PCR results showing cathepsin B (A), cathepsin S (B), cathepsin L (C), and MMP3 (D) transcript levels in 3-week-old wild type (open bars) and mutant (filled bars) RPE. Three mice of each genotype and three technical replicates per mouse were used. Values are expressed as mean ± SD, *: P<0.05; **: P<0.01 (two-tail unpaired *t*-test). **E-F.** Cathepsin B (CTSB) protein level is also increased in 3-week-old mutant RPE shown by western blot analysis (E). The results are quantified (F). Three mice of each genotype were used. Values are expressed as mean ± SD, n = 3, ***: P<0.001 (two-tail unpaired *t*-test). **G.** In the retina, mature cathepsin B is mainly in the RPE, whereas the pro-cathepsin in the neuroretina is largely unprocessed. **H-M.** Cathepsin B is localized to the apical surface of the RPE. Retinal sections from 4-week-old wild type (H-J) and mutant (K-M) were subjected to immunohistochemistry with antibodies against ezrin (H, K) and cathepsin (I, L). Merged images are shown (J, M). Retinal layers are labeled to the left. ONL: outer nuclear layer; IS: inner segment; OS: outer segment; RPE: retina pigment epithelium. Scale bar = 50μm.

Cathepsin B is a lysosomal cysteine protease normally found ubiquitously in cells and tissues [59]. In malignant tumors, the expression of cathepsin B is highly upregulated and mature cathepsin B is secreted to the cell surface where it can degrade ECM proteins. The degradation of ECM proteins by cathepsin B is required for tumor cell invasion and metastasis [53, 55, 56, 60]. To test if molecular changes of cathepsin B could be responsible for the reduced adhesion between RPE and neural retina in the *Lyst* mutant eye, cathepsin B protein levels were measured by western blot analysis. In the *Lyst* mutant eyecups, a significant elevation of mature cathepsin B protein was observed (Fig. 5E, F). Mature cathepsin B, which was mainly found in the RPE samples and not in neural retina samples, was dramatically increased relative to controls (Fig. 5G). By immunohistochemical staining, a strong cathepsin B signal was observed at the apical surface of the RPE, which colocalized with ezrin (Fig. 5H-M), and juxtaposed to the rhodopsin-labeled POS (Fig. S10). Combined with the QPCR and western blot results, these results indicate that perhaps the elevated levels of mature cathepsin B on the apical surface of RPE and interfacing POS may contribute to the reduced retinal adhesion in the *Lyst* mutant mice.

### The level of oxidative stress is increased in the *Lyst* mutant RPE

Previous studies have reported elevated expression levels of cysteine cathepsins in mouse RPE exposed to oxidative stress [58]. MMPs were also reported to be upregulated by oxidative stress in many other cell types [61–64]. These data suggest that the observed increase in levels of cysteine cathepsins and MMP3 of *Lyst* mutant RPE cells may be caused by elevated levels of oxidative stress as well. By western analysis, we determined that the level of 4-hydroxynonenal (4-HNE)-modified protein, a widely used biomarker for oxidative stress, was increased in the *Lyst* mutant RPE cells compared to wild type control (Fig. 6). Thus, we postulate that the oxidative stress due to *Lyst* deficiency results in elevation of secreted proteases in the IPM between the RPE and the photoreceptors.

**Fig. 6.**
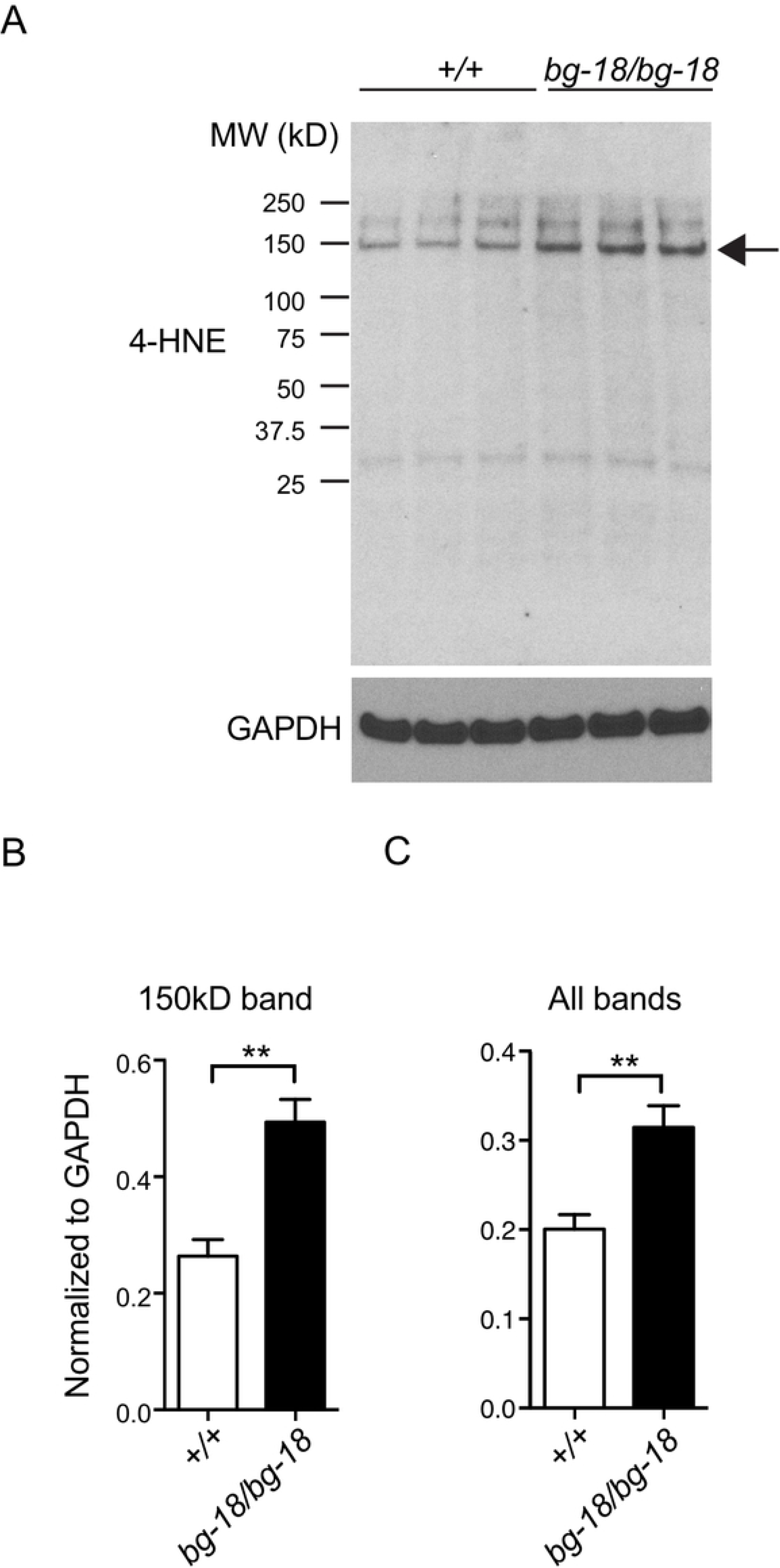
Increased oxidative stress in the *Lyst* mutant RPE. (**A**) Western blot analysis of 4-HNE modified proteins in 4-week-old wild type (+/+) and mutant (*bg-18/bg-18*) RPE. (**B-C**) The 150 kDa band on the western (A, arrow), and all bands including the 150 kDa band are quantified, respectively.

## DISCUSSION

In this study, our results demonstrate for the first time that LYST plays an important role in phagosome processing in RPE cells. We observed an abnormal accumulation of phagosomes in RPE cells in *Lyst* mutant mice (Fig. 2 and Fig. S5). Previous studies have suggested that phagocytosis of photoreceptor outer segments by RPE cells causes oxidative stress. For example, long-term daily feeding of rod OS increases the number of intracellular autofluorescent granules and increases catalase activity in cultured human RPE cells [65, 66], and rod OS uptake by cultured human RPE cells increases oxygen consumption and intracellular H_2_O_2_ production [67]. Thus, we reasoned that the accumulation of phagosomes may cause an increase in oxidative stress in the *Lyst* mutant RPE cells. In support of this hypothesis, studies have shown that slowed degradation of POS phagosomes causes oxidative stress characterized by the increase of oxidative stress markers, i.e., malondialdehyde (MDA) or 4-HNE levels, in the RPE [68, 69]. Our data shows that the levels of 4-HNE-modified proteins are increased in the *Lyst* mutant RPE compared with those in the wild type control samples, indicative of oxidative stress (Fig. 5).

Previous studies in tumor tissues suggested that oxidative stress could cause elevation of secreted cathepsin B and MMPs, which contributed to the digestion of extracellular matrix during tumor metastasis [53, 55, 56, 60]. In addition, in a mouse model of chronic oxidative stress, termed Hyperoxia-Related Retinal Degeneration (HRRD), elevation of transcript and protein levels of cathepsin B, L, and S in the RPE have been reported [58]. Strikingly, we also observed an increase in cathepsin B, L, and S, as well as MMP3 mRNA levels in *Lyst* mutant RPE (Fig. 4A-F). Our data further showed that cathepsin B is localized to the apical surface of RPE (Fig. 4H-M), where it could potentially degrade IPM proteins and thus affect the adhesion of the neuroretina to the RPE. Increased protease activity on the RPE apical surface could promote IPM remodeling and reduce adhesion of the neuroretina to the RPE. Taken together, our results suggest that in *Lyst* mutant RPE, phagosome accumulation leads to oxidative stress, which results in increased expression of proteases on the RPE apical surface, thus reducing overall RPE-retinal adhesion. Although we observed visible retinal detachment only after mechanical stress during removal of the eye from the animal, it is known that blunt ocular trauma is a common cause for retinal detachment [70–72]. Our findings suggest that Chediak-Higashi patients may be at increased risk of retinal detachment after ocular trauma.

Oxidative stress contributes to the pathogenesis of many neurodegenerative diseases including age-related macular degeneration (AMD). In addition, cathepsins have been implicated in the pathogenesis of AMD [58, 73–75]. Whether oxidative stress contributes to AMD via aberrant activation of cathepsins is still unclear. Research on the regulation of cysteine cathepsins in the context of oxidative stress may provide new therapeutic targets for AMD. Thus, the *Lyst* mutant mice may be a valuable genetic model to study the impact of chronic oxidative stress on the RPE.

RPE cells are highly phagocytic, and thus examining this cell type provides a unique opportunity to study phagocytosis *in vivo*. A similar pathological pathway described here may also exist in *Lyst*-deficient leukocytes where excessive secreted cathepsin B and other secreted proteases may cleave surface antigen or receptors, thus contributing to the decreased immune response seen in CHS patients. Alternatively, *Lyst* deficiency may primarily reduce the immune response through decreased digestion of phagosomes by lysosomes in leukocytes, which is critical for antigen presentation. The exact outcome caused by phagosome accumulation-induced oxidative stress in leukocytes versus cells in tissue such as the RPE will have to be determined experimentally in the future.

## ACKNOWLEDGMENTS

We thank the Sequencing, Histology and Imaging Sciences, and Multimedia Services at The Jackson Laboratory for their assistance in our studies. We also thank Ms. Melissa Berry for nomenclature review of the manuscript.

## Author contributions

X.J., L.Z., P.M.N, and J.K.N. designed the research study; X.J., L.Z., B.F. and J.W. performed the experiments; J.W. and B.C. contributed the mouse model and identified the *bg-18* mutation; X.J., L.Z., A.U. and D.S.W. analyzed data; and X.J., L.Z. and A.U. wrote the paper; D.S.W., J.K.N. and P.M.N. edited the manuscript.

